# Recent selection of candidate genes for mammal domestication in Europeans and language change in Europe: a hypothesis

**DOI:** 10.1101/684621

**Authors:** Antonio Benítez-Burraco, Evgeny Chekalin, Sergey Bruskin, Irina Morozova

## Abstract

Human evolution resulted from changes in our biology, behavior, and culture. One source of these changes has been hypothesized to be our self-domestication (that is, the development in humans of features commonly found in domesticated strains of mammals, seemingly as a result of selection for reduced aggression). Signals of domestication, notably brain size reduction, have increased in recent times. In this paper we compare whole-genome data between Late Neolithic/Bronze Age individuals and modern Europeans and show that genes associated with mammal domestication and with neural crest development and function are significantly differently enriched in nonsynonymous single nucleotide polymorphisms between these two groups. We hypothesize how these changes might account for the increased features of self-domestication in modern humans and ultimately, for subtle recent changes in human cognition and behavior, including language.

## Introduction

Human evolution has entailed multiple changes in our body, cognition, and behavior. These changes are expected to have resulted from selected mutations in selected genes (Grossman et al., 2013; Pääbo, 2014; Field et al., 2016) or from changes in the regulatory landscape of shared genes (Gokhman et al., 2014). Environmental factors, and particularly human culture resulting in a human-specific niche, are expected to have had an important impact on our genome too, because of the relaxation of natural selection, as well as the active selection resulting from some cultural practices (Laland et al., 2010). Beyond well-known cases mostly involving physiological adaptations (like lactase persistence, adaptation to cold climate, and adaptation to high altitude), the complex interaction between biology and culture during human evolution is poorly understood, particularly, regarding human cognition and some of its distinctive features, most notably human language. One recent hypothesis argues that many human distinctive features might have resulted from our self-domestication in response to an early selection towards increased in-group prosociality and reduced aggression (Hare, 2017; Wrangham, 2018). The parallels between domesticated animals and humans (including differences with extant primates like chimps as well as with extinct hominins) have been explored in detail by several authors (Shea, 1989; Leach, 2003; Somel et al., 2009; Zollikofer and Ponce de León, 2010; Herrmann et al., 2011; Plavcan, 2012; Stringer, 2016; Hare, 2017; Thomas and Kirby, 2018). This set of common features, impacting on the skull/brain, the face, or the skin, but also on development (paedomorphosis, reduction of sexual dimorphism) and behavior (neotenous behavior, reduced reactive aggression, increased prosocial behavior, increased play behavior), has been hypothesized to result from the hypofunction of the neural crest (NC) (Wilkins et al. 2014). Recent genomic analyses of dogs and domesticated foxes have revealed enrichments of genes linked to NC function (Pendleton et al., 2018; Wang et al., 2018). Signs of self-domestication in humans have increased in recent times (reviewed in Hare, 2017). Interestingly too, features of domestication are found abnormal (either increased or attenuated) in clinical conditions impacting on our cognitive abilities, including language, like autism spectrum disorder (Benítez-Burraco et al., 2016) or schizophrenia (Benítez-Burraco et al., 2017). At the same time, genomic regions associated with dog-human communication contain genes related to human social disorders, particularly autism spectrum disorder (Persson et al., 2016).

Not surprisingly then, self-domestication has been invoked to account for the emergence of one of the most relevant human-specific traits, namely, our cognitive ability to learn and use languages, commonly referred to as our *language-readiness* (Benítez-Burraco et al., 2018), but also of the sort of languages we use nowadays for communicating (Benítez-Burraco and Kempe, 2018; Thomas and Kirby, 2018). In a nutshell, being able to learn and use a language depends on having a brain with the proper hardware, but also on living in a cultural environment with the proper triggering stimuli. Putting this differently, our cognition accounts for many aspects of the languages we speak, but some language features are seemingly an adaptation to the physical and human-made environment, and impact in turn, more or less permanently, on our cognitive architecture. Interestingly, human self-domestication can contribute to both processes, because it gives raise to brain/cognitive changes (see Herrmann et al., 2010 for primates), but also contributes to the creation of a niche that enables the emergence of specific aspects of language complexity (like complex syntax) via a cultural mechanism, because of its enhancing impact on language learning by children, language teaching by caregivers, and language play (Benítez-Burraco and Kempe, 2018; Thomas and Kirby, 2018).

As noted above, we have detailed characterizations of the genetic differences between *Homo sapiens sapiens* (henceforth, anatomically-modern humans, AMHs) and our closest relatives, namely, Denisovans and Neanderthals (Grossman et al., 2013; Pääbo, 2014; Field et al., 2016). We also have tentative accounts of the genetic and epigenetic changes important for the emergence of our language-readiness (Boeckx and Benítez-Burraco, 2014a), as well as some preliminary hypotheses about how these changes could have been translated to changes in the sort of cognitive abilities that are needed for acquiring and mastering a language (Murphy and Benítez-Burraco 2018). Interestingly, one recent genetic research has shown that candidate genes for domestication in mammals are overrepresented among the genes under positive selection in AMHs compared to extinct hominins (Theofanopoulou et al., 2017). However, no evidence of when these changes were selected is available. Moreover, because of the attested effect of environmental factors, and more generally, our mode of life, on our morphology, physiology, and behavior, as noted above, it is not clear either whether the observed differences between early AMHs and present-day AMHs resulted from the enhancement of our self-domestication, or are instead an unrelated consequence of our adaptation to new, human-made environments.

Overall, because self-domestication can be considered a process with different degrees of completion, and because of our ignorance of the timing of the self-domestication event(s), we regard of interest to check, specifically, whether genomic signals of domestication have intensified recently in AMHs. If this was the case, one could argue that some late changes in human evolution with an impact on language are certainly associated with our self-domestication, rather than simply with changes of life. In a recent paper (Chekalin et al., 2019), we compared whole-genome data between Late Neolithic/Bronze Age individuals from 6000 years ago and modern Europeans and found that several biological pathways are significantly differently enriched in nonsynonymous single nucleotide polymorphisms (SNPs) in these two groups. We then argued that these changes, with an impact on metabolism, immune response, physical behavior, perception, reproduction, and cognition, could have been triggered and shaped by cultural practices, particularly, by important changes occurred in Europe at that age. In this paper, we have relied on the same genomic data and asked whether a genetic signature of enhanced self-domestication can be found that accounts for the attested enhancement of domestication features in late AMHs and seemingly also, for some of the recent changes we attested in our previous work. To answer this question, we have analyzed the same two samples of Europeans (Late Neolithic/Bronze Age and modern ones), in order to compare the numbers of nonsynonymous mutations in the groups of genes associated with self-domestication and NC development and function. Because our focus of interest is put on language, we have also interrogated whether robust candidates for language development and evolution show signals of recent selection in our European samples that can account for the hypothesized recent changes in their languages, or whether these changes can be confidently assigned to recent selection of candidates for domestication.

## Materials and Methods

Regarding the samples of Europeans, we compared whole genome data from 150 ancient samples dated between 3,500 and 1,000 BCE (Gamba et al., 2014; Allentoft et al., 2015; Haak et al., 2015; Mathieson et al., 2015) with data on 305 modern Europeans genotyped in the framework of the 1,000 Genomes Project (Genomes Project et al., 2015), under the assumption that modern Europeans are genetic descendants of the Bronze Age Europeans, as described and discussed in detail in Chekalin et al. (2019) (Figure 1). To minimize the influence of demographic events, we restricted our ancient data to individuals attributed to Yamnaya and related archaeological cultures whose representatives probably spoke Indo-European languages (Gamba, et al. 2014; Allentoft, et al. 2015; Haak, et al. 2015; Mathieson, et al. 2015), and our modern data by European carriers of Indo-European languages, which are considered to be the descendents of Yamnaya individuals. The genetic continuity between the ancient and the modern groups was confirmed using Principal component and reAdmix analysis (Chekalin et al., 2019).

**Figure 1.**
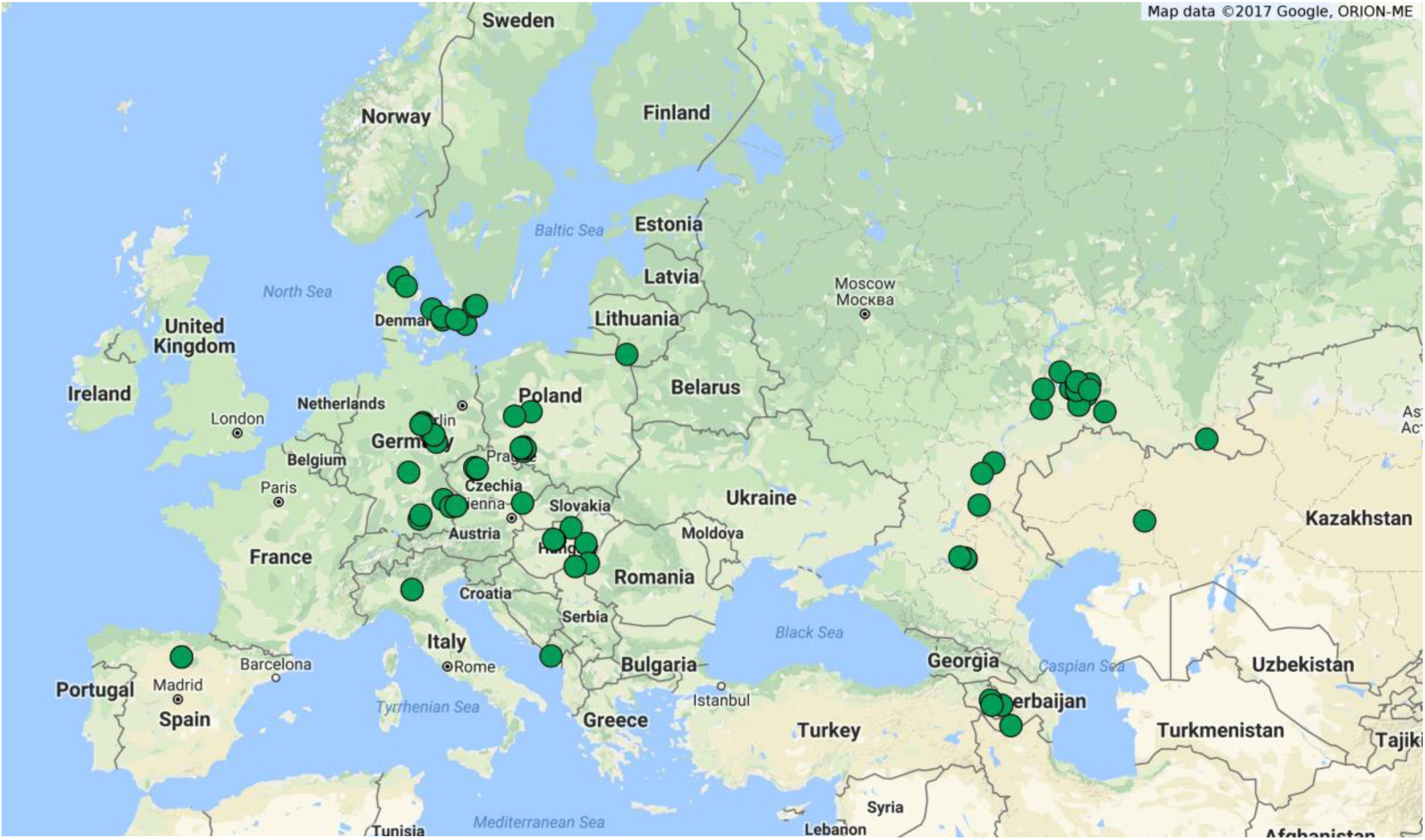
Location of ancient samples analyzed in the study.

Regarding genes potentially related to self-domestication, we considered four groups of genes. Our first set encompassed 764 candidate genes for mammal domestication (Supplemental table 1; column A). This list resulted from merging genes that have been found positively selected in several domesticates, including zebra fish, quail, chicken, Guinea pig, pig, rat, dog, cat, cattle, domesticated fox, horse, rabbit, and sheep (Womack et al., 2005; Trut et al., 2009; Albert et al., 2011; Axelsson et al., 2013; Bellone et al., 2013; Carneiro et al., 2014; Montague et al., 2014; Qanbari et al., 2014; Schubert et al., 2014; Wilkins et al., 2014; Wright et al., 2015; Cagan et al., 2016; Freedman et al., 2016; Zapata et al., 2016; Benítez-Burraco et al., 2017; Theofanopoulou et al., 2017; Pendleton et al., 2018). A second list encompassed the 41 candidates for domestication highlighted by Theofanopoulou and collaborators (2017) as showing evidence of positive selection in AMHs compared to Neanderthals/Denisovans (Supplemental table 1; column B). In view of the suggested role of the NC in the emergence of features of domestication, we considered as well genes important for NC development and function, which are the ones we compiled for our paper on domestication and schizophrenia (Benítez-Burraco et al., 2017). This list encompassed 89 genes (Supplemental table 1; column C), which we gathered using pathogenic and functional criteria: neurochristopathy-associated genes annotated in the OMIM database (http://omim.org/), NC markers, genes that are functionally involved in NC induction and specification, genes involved in NC signaling (within NC-derived structures), and genes involved in cranial NC differentiation. Finally, we considered as well the “core” genes (n= 16) highlighted by Wilkins and collaborators (2014) as key candidates for the “domestication syndrome” in mammals (Supplemental table 1; column D). Because these four sets of genes are involved in domestication and/or NC development and function, we expected that they help to characterize recent changes in human behavior and cognition resulting from self-domestication with an impact on language abilities and structure: although direct causal links between some of our candidates and domestication traits of interest can be regarded hard to find, current approaches to the genetics of complex traits based on genome-wide association studies (GWAS), of the sort we have relied on for compiling our lists of candidates, make this endeavor feasible (Goddard et al., 2016, Guo et al., 2018).

Finally, regarding the core candidates for language development and evolution, we have made use of two sets of genes. The first list encompassed 50 robust candidates for the two main specific language disorders: developmental dyslexia (DD) and specific language disorder (SLI) (Supplemental table 1; column E). These are cognitive disorders impacting on language only, either on phonology mostly (DD) or on grammar mostly (SLI). For DD, we have relied on the updated list of candidates for this condition provided by Paracchini et al. (2016), which includes genes resulting from candidate association studies, GWAs, quantitative GWAs, copy number variation (CNV) studies, and next-generation sequencing (NGS) analyses; we also surveyed the literature via PubMed (ncbi.nlm.nih.gov/pubmed), looking for additional candidate genes. Concerning SLI, we relied on the recent list of candidates compiled by Chen et al. (2017), but we also considered the literature survey and the results by Pettigrew et al. (2016), who compiled strong candidates resulting from linkage analyses, GWA studies, and NGS analyses. Similarly to what we did for DD, we also surveyed the literature via PubMed looking for additional robust candidates for this condition. The second list encompassed 152 genes (Supplemental table 1; column F). It contained strong candidates for language evolution, as compiled by Boeckx and Benítez-Burraco (2014a; 2014b) and Benítez-Burraco and Boeckx (2015). These genes are primarily expected to account for the globularization of our skull/brain and for the cognitive changes resulting in our language-readiness. As discussed in detail in these three papers, these 152 genes fulfil several of the following criteria: i) they have changed (and/or interact with genes that have changed) after our separation from extinct hominins, this including changes in their coding regions and/or their regulation; ii) they are involved in brain development, wiring, and/or function; and/or iii) they are candidates for language deficits in broader, highly prevalent cognitive disorders, particularly, autism spectrum disorder and schizophrenia (see Benítez-Burraco and Murphy, 2016; Murphy and Benítez-Burraco, 2016; 2017 for additional details about their role of many of these genes in language processing).

We wish to note that despite the fact that it is widely accepted that mutations in cis-regulatory regions play a very important role in evolution (King and Wilson, 1975 and many others), the functions of most of the SNPs in the regulatory regions are not yet known, and no confident database of these sort of changes in the human lineage is currently available. For this reason, our analysis was restricted by genome coding regions only. For each of the six sets of genes, we performed the calculations analogous to those done in the paper of Chekalin and colleagues (2019). Briefly, we calculated the significance of the differences in the counts of nonsynonymous SNPs in the sets between ancient and modern Europeans. The significance was assessed via differential non-synonymous SNP enrichment (DNSE) scores using the following formula:

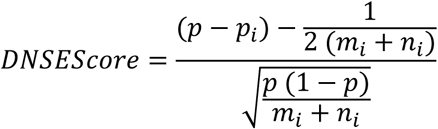

where *p* is the fraction of ancient non-synonymous SNPs in the whole analyzed set, *n* is the amount of ancient nonsynonymous SNPs in the analyzed sets of genes, m is the amount of modern nonsynonymous SNPs in the analyzed sets of genes. The fraction *p*_*i*_ of ancient non-synonymous SNPs in the *i*^th^ set is *p*_*i*_ = *n*_*i*_/(*n*_*i*_ + *m*_*i*_), *w*here *n*_*i*_ is the amount of ancient non-synonymous SNPs in the *i*^th^ set, *m*_*i*_ is amount of modern nonsynonymous SNPs in the *i*^th^ set. From acquired numbers enrichment DNSE scores were computed for every pathway with continuity correction (Fleiss, et al., 2003). We assume that during neutral evolution similar sets of genes accumulate nonsynonymous SNPs at the same rate in both ancient and modern groups, and during enrichment analysis such sets would fit a normal distribution, while sets that are affected by evolutionary pressure would be outliers from this distribution. For calculated DNSE scores (distributed normally, Shapiro-Wilk test *p-*value > 0.01), *p*-values were calculated using Bonferroni and Benjamini-Hochberg corrections. The gene sets were considered to be differentially enriched if absolute value of the DNSE score > 4, and the adjusted *p-*value < 0.01. Negative DNSE values indicated accumulation of mutations in genomes of modern Europeans in comparison with ancient Europeans, while positive score values indicated an opposite pattern.

To make sure that the observed differences in DNSE scores between ancient and modern groups are not caused by ancient DNA artefacts, we also calculated differential synonymous SNP enrichment (DSSE) scores using the same formula as for DNSE and then normalize nonsynonymous SNPs on synonymous SNPs (Chekalin et al., 2019). To validate our method, we compared it with the method implemented by Somel et al. (2013). Calculation of DNSE scores for two groups analyzed in Somel et al. (2013), chimpanzee and their ancestors, revealed the same results as in the paper of Somel and colleagues (Chekalin et al., 2019). Additional tests, including the Spearman’s correlation coefficient between enrichment score and fraction of covered length and analysis of dependence of enrichment on coverage and gene size, confirmed that revealed differences are not the results of ancient DNA features (Chekalin et al., 2019).

## Results

As expected, we found no significant differences between groups in synonymous SNP enrichment (Table 1), taking into account the neutral character of these mutations. At the same time, we found a significant difference in enrichment in nonsynonymous SNPs between the Bronze Age and present-day European individuals. Specifically, we found that candidates for domestication have been accumulating nonsynonymous mutations during the past 6,000 years, whereas candidates for NC exhibit fewer nonsynonymous mutations in present-day humans than in Bronze Age humans. By the reasons we provide in our 2019 paper (Chekalin et al., 2019), these differences are not expected to be caused by an insufficient sequence coverage of Bronze Age individuals or by general inter-population differences between the two groups. By contrast, we found no significant selection signals in the group of domestication candidates positively selected after our split from Neanderthals and Denisovans, nor in “core” candidates for the domestication syndrome (Table 1). We have found no signals of selection either in genes related to language (neither in candidate genes for language disorders, nor in genes related to language evolution). We discuss now these results and argue that this lack of selection reinforces our hypothesis that recent changes in language features can be preferentially linked to our enhanced self-domestication.

**Table 1.** Differently enriched sets of genes in ancient and modern groups. Positive enrichment score values correspond to pathways that have more SNPs in genomes of ancient individuals, while negative DNSE values correspond to pathways that have more SNPs in genomes of modern Europeans. AMH refers to anatomically-modern humans (compared to Neanderthals and Denisovans)

| Set of genes | Ancient SNPs count | Modern SNPs count | Enrichment score | p-value | p-value adjusted Bonferroni | Enrichment |
| --- | --- | --- | --- | --- | --- | --- |
| Synonymous SNPs |  |  |  |  |  |  |
| Domestication | 2165 | 3759 | 1.24 | 0.2138 | 1 | No |
| Neural crest | 194 | 315 | 1.19 | 0.2353 | 1 | No |
| Domestication syndrome | 49 | 76 | 0.98 | 0.326 | 1 | No |
| Positive selection in AMH | 144 | 268 | -0.25 | 0.8012 | 1 | No |
| Language disorders | 110 | 177 | 1.02 | 0.3057 | 1 | No |
| Language evolution | 462 | 837 | -0.10 | 0.9193 | 1 | No |
| Nonsynonymous SNPs |  |  |  |  |  |  |
| Domestication | 2440 | 4844 | -4.32 | $1.58 \times 10^{-5}$ | 0.005 | Modern |
| Neural crest | 213 | 240 | 5.01 | $5.36 \times 10^{-7}$ | $2 \times 10^{-4}$ | Ancient |
| Domestication syndrome | 61 | 73 | 2.49 | 0.0127 | 1 | No |
| Positive selection in AMH | 138 | 231 | 0.69 | 0.4885 | 1 | No |
| Language disorders | 119 | 241 | -0.90 | 0,3694 | 1 | No |
| Language evolution | 532 | 871 | 1.87 | 0,0614 | 1 | No |

## Discussion

Self-domestication has been claimed to account for key aspects of human evolution, including the creation of the cultural niche that allowed complex languages to emerge. Although signals of domestication have seemingly increased recently, as showed by the paleoanthropological record (Leach 2003; Zollikofer and Ponce de León, 2010; Stringer, 2016), it is not clear if they specifically resulted from genomic changes, that incidentally enable to provide a more precise chronological account of the self-domestication events, as it has been possible with several domesticated mammal species (Driscoll et al., 2007; Nomura et al., 2013; Orlando et al., 2013; Freedman et al, 2014; Qiu et al., 2015; Botigué et al., 2017). At present, only one study has addressed this issue, concluding that statistically significant overlaps exist between selective sweep screens in AMHs and several domesticated species (Theofanopoulou et al., 2017). Nonetheless, this study is inconclusive about the timing of the self-domestication events, as it relied on previously published (but limited) comparisons between AMHs and Neanderthals and Denisovans by Prüfer et al. (2014), Racimo (2016) and Peyrégne et al. (2017).

In this paper, we have shown that genes associated with self-domestication and with NC development and function have been selected during the last 6,000 years in Europe, a period when important changes in human behavior and culture occurred, including the spread of agricultural practices and sedentism, urbanization, increasing in population density, development of trading routs, globalization, etc. These changes reshaped not only the gene pool of Europe, but also modified its linguistic landscape, because Neolithic languages were almost totally replaced by Indo-European languages (Bouckaert et al., 2012; de Barros et al., 2018; Mathieson, 2018, among many others).

The group of genes that are candidates for domestication in mammals has demonstrated the accumulation of nonsynonymous mutations in the genomes of present-day Europeans in comparison to Bronze Age ones. This group consists of 764 genes which can be responsible for a number of different biological processes. Further detailed study of this group with its subsequent division into smaller subsets will probably allow us to reveal more diverse patterns of selections for these genes. Likewise, we have found the decrease in nonsynonymous mutations in the modern group in comparison to ancient Europeans in the candidate genes for NC development and function. This can be the evidence of negative or, on the opposite, strong positive selection in the genes responsible for development and function of NC. This finding seemingly reinforces the suggested role of changes in NC function as the main trigger of domestication (and self-domestication) events.

As far as our cognitive evolution is concerned, enhanced self-domestication has been recently claimed to contribute to the transition from the so-called esoteric languages, typically spoken by close-knit, small human communities that share a considerable amount of knowledge about the environment, to exoteric languages, better designed for communicating decontextualized information to strangers (Benítez-Burraco and Kempe, 2018). In brief, self-domestication seemingly resulted in a less aggressive behaviour that facilitated the establishment of larger and more complex social networks and enhanced contacts with strangers, which are factors that favour the emergence of exoteric features in languages (phonological simplification, morphological transparency and regularity, expanded vocabularies, and more complex syntax). As recently suggested by Sánchez-Villagra and van Schaik (2019), some of the physical features brought about by self-domestication, like our childish face, might have contributed to these social changes, acting as signals of friendliness and social tolerance. Ultimately, the extended juvenile period also resulting from self-domestication might have potentiated language learning by children and language teaching by caregivers, enabling the mastering of exoteric languages, which are most costly to process and learn (Benítez-Burraco and Kempe, 2018). Our hypothesis here is that for the reasons mentioned above, in Europe (and seemingly in other parts, but this needs to be checked) this transition to exotericity could be linked to the increased domestication features found among Europeans in that period. Importantly, exoteric languages are expected to demand some cognitive adaptation, because their more complex syntax and expanded vocabularies need more working memory capacity, more executive control, and more declarative knowledge to be learnt and mastered (see Benítez-Burraco and Kempe, 2018 for a detailed discussion). Interestingly, in our previous work we found in our European samples evidence of selection of two pathways related to cognition, particularly, to long-term potentiation and dopaminergic synapse: whereas the former underlies synaptic plasticity and ultimately, memory and learning abilities, the latter is involved in executive control (Chekalin et al., 2019). Importantly, these two pathways do not have any shared genes with NC genes. Therefore, the common pattern found for these two groups (i.e. decrease in nonsynonymous mutations during the last 6,000 years) is seemingly due to not shared genetic background but, probably, to common external factors. In our 2019 paper (Chekalin et al., 2019) we suggested that this selection might be related to changes in ways of information presentation, perception, and transmission. Now, we hypothesize that it might be (also) related to the transition from esoteric to exoteric languages in Europe via enhanced self-domestication. The feasibility of this hypothesis is suggested by the fact that we have found no signals of selection in genes related to language disorders or language evolution. This lack of selection reinforces the view that if gene-culture coevolution has played some role in the recent changes attested in language features in Europe, this has not involved the genes that contributed to the emergence of our language-readiness and of modern speech, but genes related to domestication, that might have favored the cultural evolution of languages because of their impact on our behavior and cognition.

Likewise, we have found no signals of recent selection in Europeans in “core” candidates for domestication (Wilkins et al. 2014) or in candidates for domestication showing signals of positive selection in AMHs compared to Neanderthals and Denisovans (Theofanopoulou et al., 2017). This suggests that these two sets of genes might have been selected earlier in our history, plausibly accounting for the domesticated phenotype already exhibited by our ancestors well before 6.000 years ago (e.g. Cieri et al., 2014). By contrast, more recent self-domestication events, like the sort we highlight here in connection with the transition to exoteric-like languages, seemingly resulted from selection in other candidates for domestication and NC development and function.

Overall, our results suggest that human self-domestication is an ongoing process. Accordingly, we can expect that it contributed to past and recent changes in the human body, behavior, and culture, and seemingly cognition, with a potential impact on language evolution. Selection on different sets of candidates for domestication seems to account for the different stages of the human self-domestication process. Ultimately, we can suggest that domestication and cultural practices are interdependent; they trigger each other during the ongoing cultural and biological evolution of human beings.

## Acknowledgements

This research was funded by the Spanish Ministry of Economy and Competitiveness (grant FFI2016-78034-C2-2-P [AEI/FEDER, UE] to ABB). We thank Adam Wilkins for some valuable commentaries on a previous version of this manuscript.

## Author contribution statement

ABB conceived the paper. ABB and IM wrote the manuscript. EC, SB, and IM performed the analyses and discussed the results.

